# Patterns of socio-cognitive stratification and perinatal risk in the child brain

**DOI:** 10.1101/839969

**Authors:** Dag Alnæs, Tobias Kaufmann, Andre F. Marquand, Stephen M. Smith, Lars T. Westlye

## Abstract

The expanding behavioral repertoire of the developing brain during childhood and adolescence is shaped by complex brain-environment interactions and flavored by unique life experiences. The transition into young adulthood offer opportunities for adaptation and growth, but also increased susceptibility to environmental perturbations, such as the characteristics of social relationships, family environment, quality of schools and activities, financial security, urbanization and pollution, drugs, cultural practices, and values, that all act in concert with our genetic architecture and biology. Our multivariate brain-behavior mapping in 7,577 children aged 9-11 years across 585 brain imaging phenotypes, and 617 cognitive, behavioral, psychosocial and socioeconomic measures revealed three population modes of brain co-variation, which were robust as assessed by cross-validation and permutation testing, taking into account siblings and twins, identified using genetic data. The first mode revealed traces of perinatal complications, including pre-term and twin-birth, eclampsia and toxemia, shorter period of breast feeding and lower cognitive scores, with higher cortical thickness and lower cortical areas and volumes. The second mode reflected a pattern of socio-cognitive stratification, linking lower cognitive ability and socioeconomic status to lower cortical thickness, area and volumes. The third mote captured a pattern related to urbanicity, with particulate matter pollution (PM^25^) inversely related to home value, walkability and population density, associated with diffusion properties of white matter tracts. These results underscore the importance of a multidimensional and interdisciplinary understanding, integrating social, psychological and biological sciences, to map the constituents of healthy development and to identify factors that may precede maladjustment and mental illness.

## Introduction

The complexity and idiosyncratic characteristics of the human mind originates in an intricate web of interactions between genes, brain circuits, behaviors, economic, social and cultural factors during childhood and adolescence. The major life changes associated with the transition into young adulthood offer opportunities for adaptation and growth, but also increased susceptibility to detrimental perturbations, such as the characteristics of social and parental relationships, family environment, quality of schools and activities, economic security, urbanization and pollution, drugs, cultural practices, and values, that all act in concert with our genetic architecture and biology. A multidimensional understanding of the interplay of these factors is paramount to identify the constituents of healthy development and to identify factors that may precede maladjustment and mental illness.

Population-based neuroimaging now allows us to take a birds-eye view on this stupendous multiplicity, and to bring hitherto unseen patterns into focus^1^. The Adolescent Brain Cognitive Development (ABCD) study^2^ provides brain images of more than 10,000 children aged 9-11 years across the US and includes a broad range of cognitive, behavioral, clinical, psychosocial and socioeconomic measures. While each neuroimaging feature typically explains a minute amount of unique variance in behavioural outcome^3, 4^, their combined predictive value is non-negligible, including predictive patterns for identification of individuals^5, 6^ and characteristics such as age^7, 8^, cognitive ability^9^ and psychopathology^9^. This added value of multivariate and combinatorial approaches for prediction of complex traits is highly analogous to the substantial polygenic accumulation of small effects in the genetic architecture of complex human traits and disorders^10, 11^.

Adolescence is a transition period between childhood to adulthood and a period of protracted brain maturation, associated with heightened sensitivity to the social and cultural environment^12^. For most individuals, this transition results in successful acquirement of skills and coping strategies required for adulthood and subsequent independence from caregivers, however it is also period of increased risk for mental health issues^13^, with possible life-long repercussions. Mapping positive and negative factors impacting the brain as well as psychological adjustment during the transition from childhood to adulthood is therefore of pivotal importance. Combining levels of information using latent-variable approaches which model all available information may reveal interpretable patterns among multiple brain imaging features and variables such as cognition and socio-demographics^4, 14^. One recent example revealed that a wide range of cognitive, clinical and lifestyle measures constitute a “positive-negative” dimension associated with adult brain network functional connectivity^1^.

Here we used an analogous approach in 7,577 children aged 9-11 from the ABCD-study, collected across 21 sites across the US, combining canonical correlation analysis (CCA) with independent component analysis (ICA) to derive population-level modes of co-variation, linking behavioral, psychosocial, socioeconomic and demographical variables (behavioral measures) to a wide set of neuroimaging phenotypes. Each resulting mode represents an association between a linear combination of behavioral measures with a separate combination of imaging features that show similar variation across participants^14^. In order to avoid overfitting, which is particularly important when employing data-driven approaches, and due to the high number of inter-correlated features, CCA was performed after data reduction with principal component analysis, and robustness and reliability of the identified modes were assessed using stratified cross-validation and permutation testing with restricted exchangeability, taking into account siblings and twins based on participant’s genetic data. To express results in the original variable-space, CCA-ICA subject weights were correlated back into the original data. Based on earlier reports of population level associations between measures of life-outcomes and brain connectivity^1^ and structure^15^ in adults, the known and rising socioeconomic inequalities in the US^16^, as well the impact of socioeconomic factors on child brain development^17, 18^, we expected to find traces of social stratification in the child brain.

## Methods

### ABCD data access

We accessed MRI, behavioral, clinical and genetic data from ABCD Annual curated release 2.0.1. The data as well as release notes including documentation of measures, scanning protocols and imaging QC can be accessed using the following NIMH data archive DOI: http://dx.doi.org/10.15154/1503209.

### Behavioral, clinical, cognitive and demographical data

Tabulated data was imported and processed using R (https://cran.r-project.org). We accessed data from 11,853 participants. **Supplementary Table 1** lists the behavioral measures included the analysis. We used the function ‘nearZeroVar’ from the R-package ‘caret’ (v. 6.0-81, https://github.com/topepo/caret/) to identify and exclude any continuous variables with zero or near-zero variance, and categorical variables with a ratio of > .95 for the most common compared to the second most common response. For each remaining variable we derived robust z-scores by calculating each scores absolute deviation from the median absolute deviation^19^ (MAD), and removed values with a z > 4 (4 x MAD). Those with a z > 3 were manually inspected: e.g. measures of facility income, time spent on phone and several measures of area deprivation have scores with z>3, but were kept in the analysis. We then excluded variables with less than 90% of retained datapoints, before excluding subjects with less than 90% retained data across the retained variables. The remaining subjects (n=11,809) were included for further analysis.

### MRI Imaging derived phenotypes

We accessed T1 and T2 (n=11,534) and DWI (n=11,400) tabulated data from ABCD curated release 2.0.1. **Supplementary Table 2** lists the MRI features included in the analysis. We included participants which passed quality assurance using the recommended QC parameters (T1: n=11,359, T2: n=10,476, DWI: n=10,414) described in the ABCD 2.0.1 Imaging Instruments Release Notes and whom had all included modalities available (n=9,811). ABCD preprocessing and QC steps are described in detail in the methodological reference for the ABCD Study by Hagler et al^20^. For each included imaging phenotype, we calculated the median absolute deviation (MAD) for each score, and removed values with MAD > 3. Subjects with less than 90% of features retained in any of the imaging modalities, and features with less than 90% of retained subjects were excluded from analysis. The remaining subjects (n=9,016) were included for further analysis.

### Genetic data

We accessed genetic data for 10,627 participants to identify siblings and twins. We used genome-wide complex trait analysis^21^ to create a genetic relationship matrix after performing the following filtering: removal of SNPs in the major histocompatibility complex (25:35 Mb region on chr6) and the inversion region of chr8 (7:13 Mb); SNPs with genotyping rate <99%, minor allele frequency < 5%, pairwise pruning of SNPs in linkage disequilibrium (r^2^ > 0.2, window of 5,000, step of 500). To account for sibling and twins in the dataset, three groups were created based on the following genetic relatedness cut-offs, <.4, >.4 & <.6, and >.8, with the two latter groups containing pairs of siblings, and used for stratified cross-validation and creation of permutation exchangeability blocks.

### Canonical correlation analysis

We performed CCA^22^ using MATLAB R2019b. Participants with MRI, behavioral and genetic data (n=7,577) were included. We applied a rank-based normal transformation to the behavioural/clinical data using ‘palm_inormal’ from FSLs PALM^23^ (v. 0.52, https://fsl.fmrib.ox.ac.uk/fsl/fslwiki/PALM). Next, we residualized all measures with respect to age and sex using linear models. Imaging phenotypes were also residualized for site/scanner, and volumetric features were also corrected for estimated total intracranial volume (eTIV from Freesurfer). eTIV was also included as variable in the analysis to capture associations with global volume, in addition to the eTIV-corrected volumes capturing associations with regional specificity. For both MRI and behavioral measures, missing values were imputed with ‘knnimpute’, replacing missing data based on the *k* nearest-neighbour columns based on Euclidian distance (*k*=3). An alternative approach without imputation is described below and did not change results. Data were then z-normalized and submitted (separately for imaging and behavioural data) to PCA (**Supplementary Fig. 1**), to avoid issues with rank deficiency and to increase robustness of estimated modes by avoiding fitting to noise. We extracted the first 200 components for both the imaging and behavioural data, and submitted these to CCA.

### Cross-validation

To assess the reliability and generalizability of the resulting CCA-modes we performed the following 10-fold cross-validation procedure: For each iteration (n=100) of the cross-validation loop the dataset was randomly divided into 10 folds, stratified by the genetic relatedness groups, and ensuring that sibling and twin pairs were kept together to avoid training on one sibling/twin in a pair, and test on the other. While keeping each fold (10% of participants) out once we submitted the remaining data (90% of participants) to PCA (separately for imaging and MRI data) and then to CCA. Next we multiplied the kept-out behavioral measure and MRI feature matrices with the estimated PCA coefficient matrices, before multiplying the resulting PCA scores with the canonical coefficients and then correlated the resulting CCA scores. Finally, we took the average of these canonical correlations across the 10 folds (**Supplementary Fig. 2**). This procedure was repeated 100 times to derive mean canonical correlations for kept-out data, and used for calculating p-values after permutation testing. We also correlated the CCA subject measure and MRI coefficients derived for kept-out participants, with those from the full analysis.

### Permutation testing

To assess significance of the resulting CCA-modes, we ran 1000 iterations of the same 10-fold cross-validation procedure described above, but with the order of participants of the imaging phenotype matrix randomly permuted in each iteration, respecting twins/sibling relationships, and collecting canonical correlations for the kept rather than the kept-out data to account for overfitting by the CCA. We then collected the maximum canonical correlation across CCA-modes (i.e. mode 1) for each permutation to form a null-distribution to calculate familywise error corrected (FWE) p-values. P-values for each of the CCA-modes were calculated by dividing the count of permuted maximum R-values (including the observed value) >= the mean of cross-validated R-values by the number of permutations. CCA-modes with a corrected p-value < .01 was included for further analysis (**Supplementary Fig. 2**).

### CCA-ICA

The canonical variates becomes increasingly difficult to interpret due to their orthogonality. Since we had more than one significant mode, and following procedures described by Miller et al^14^, we used ICA to obtain more interpretable modes: we extracted and combined the behavioral and MRI CCA-scores for the three significant variates, correlated these with the original data matrix, transformed the correlations using a Fisher Z-transform, and submitted these to ICA. We estimated three components (the number of significant and extracted CCA covariance-modes) using fastICA^24^. To assess the reliability and generalizability of the ICA decomposition we reran 100 iterations of the 10-fold cross-validation procedure described above, this time including ICA estimation after the PCA and CCA step, and then correlated ICA subject-weights derived from kept-out data to those from the full analysis (**Supplementary Fig. 2**). To assess and plot the significant CCA-ICA modes in the full original variable space, we correlated the subject weights for each CCA-ICA mode with the original age- and sex (+ eTIV) adjusted matrices. For each significant mode of population covariation, we also plotted the variable text/descriptions for the 35 variables with the highest explained variance in the original adjusted data (lists of all variables and associated descriptions, correlations and ICA weights can be found in **Supplementary Tables 3-5**). ICA subject weight histograms are shown in **Supplementary Fig. 3**. The explained variance of single variables ranged between 10% to 40% for the most highly involved items on these population modes, which is in a similar range as reported employing a similar approach in the adult UK Biobank sample^14^. For visualization purposes, we produced scatter-plots using the highest-loading variables for each mode, color-coded by each individual’s score on the respective modes (**Supplementary Fig. 4**).

### Consistency across sex and race/ethnicity

To assess the degree of similarity of the patterns across the sexes, we split the CCA-ICA subject weights by sex (**Supplementary Figs. 5–7**) and compared sex-specific subject-weight-with-variable correlations to those estimated for the full analysis. Correlations for the three modes ranged between r=.95 and r=1. The aim of this work was not to make comparisons of population sub-groups, but to detect general population patterns. Since many of the included indicators relating to inequality and socioeconomics are known to differ between ethnic minority groups, we did not regress these variables out of the data. To show that the detected patterns are generalizable we computed subject-weight-with-variable correlations for groups based on parent-ascribed race/ethnicity (**Supplementary Fig. 8–10**) excluding those with a frequency < 5% of the total sample (retaining “black”, “white”, “other”, and compared these to the full analysis. Correlations for the three modes ranged between r=.76 and r=1. These results indicate that the patterns are generalizable across sexes and ethnicity/race.

### Consistency across sites

To further assess the generalizability and robustness of the CCA-ICA patterns we computed site/scanner-wise CCA-ICA variable-correlations, and performed correlations comparing these to the full model (**Supplementary Fig. 11–13**). The pattern of the three modes are mostly consistent across sites, but with some sites deviating more from the full analysis modes than others (r=.85 - r=.22). All imaging phenotypes were adjusted for site, however, ABCD collects data at 21 sites across the continental US (https://abcdstudy.org/about) and population-level demographical differences are expected.

### Alternative approach without imputation

To ensure that the results were not affected by the imputation procedure for missing data points, we also used an alternative and previously described approach^1^ in which we estimated the subject x subject covariance matrix, ignoring missing values, before projecting this approximated covariance matrix to the nearest positive-definite covariance matrix using the MATLAB tool ‘nearestSPD’ (https://www.mathworks.com/matlabcentral/fileexchange/42885-nearestspd), thereby avoiding the need for imputation of missing values. The correlations between the CCA scores for the first three modes between the original analysis using imputation and this approach were r= .99, r=.96 and r=.96, respectively.

### Alternative number of PCA components

To investigate the impact of choosing a stricter criterion of inclusion of PCA we reran the analysis with 100 PCA components, and compared the resulting CCA scores for the first three modes with those from the original analysis, yielding correlations of r=.98, r=.91, and r=.91, respectively.

### Adjusting data for age^2^

To address the possibility of non-linear relationships between age and the various demographic, clinical and MRI measures features we reran analysis with age^2^ added along with the original confound variables, and compared the resulting CCA-ICA subject weights for the first three modes with those from the original analysis, yielding correlations of r=.99, r=.96 and r=.97, respectively.

## Results

We identified three distinct modes of co-variation, linking brain-features to perinatal and early life events, socio-cognitive factors and urbanicity (**Fig. 1**). Canonical correlations for the first three modes where were significant and robust as assessed by 10-fold cross-validation and permutation (out-of-sample *r=.61, r=.42, r=.38*, all *permuted-p = 0.001* respectively, **Supplementary Fig. 2**).

**Fig. 1:**
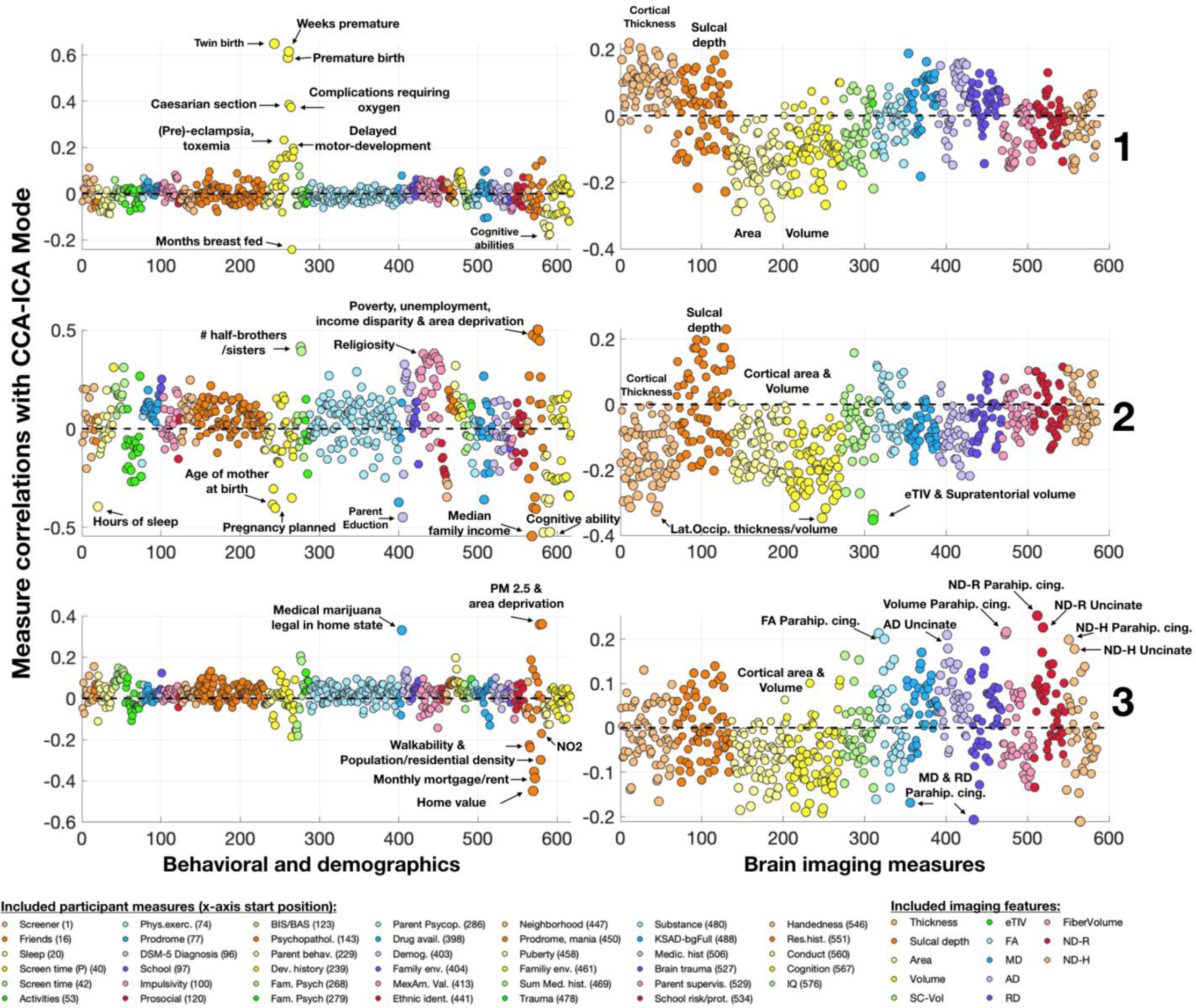
Each row represents a CCA-ICA mode. Each mode represents an association between a linear combination of behavioral measures (left) with a separate combination of imaging features (right). X-axes represent the numbered behavioral measures / imaging features. Behavioral measures legend shows starting location on x-axis for each measure (in parenthesis). Y-axes shows the correlation between each included variable with the CCA-ICA subject weights.

Mode 1 links perinatal factors and obstetric complications to cognitive ability and brain morphology in late childhood (**Fig. 2**). Having a twin, premature birth, birth complications requiring oxygen, Caesarian section, (pre)-eclampsia, toxemia and jaundice is associated with shorter duration of breast feeding, parent reported delayed motor development, lower cognitive scores and linked to a pattern of cortical morphometry and white matter diffusion measures in several brain regions, with lower cortical volume and area and higher thickness in middle temporal, lateral orbitofrontal and inferior parietal cortex among the highest-loading imaging features.

**Fig. 2:**
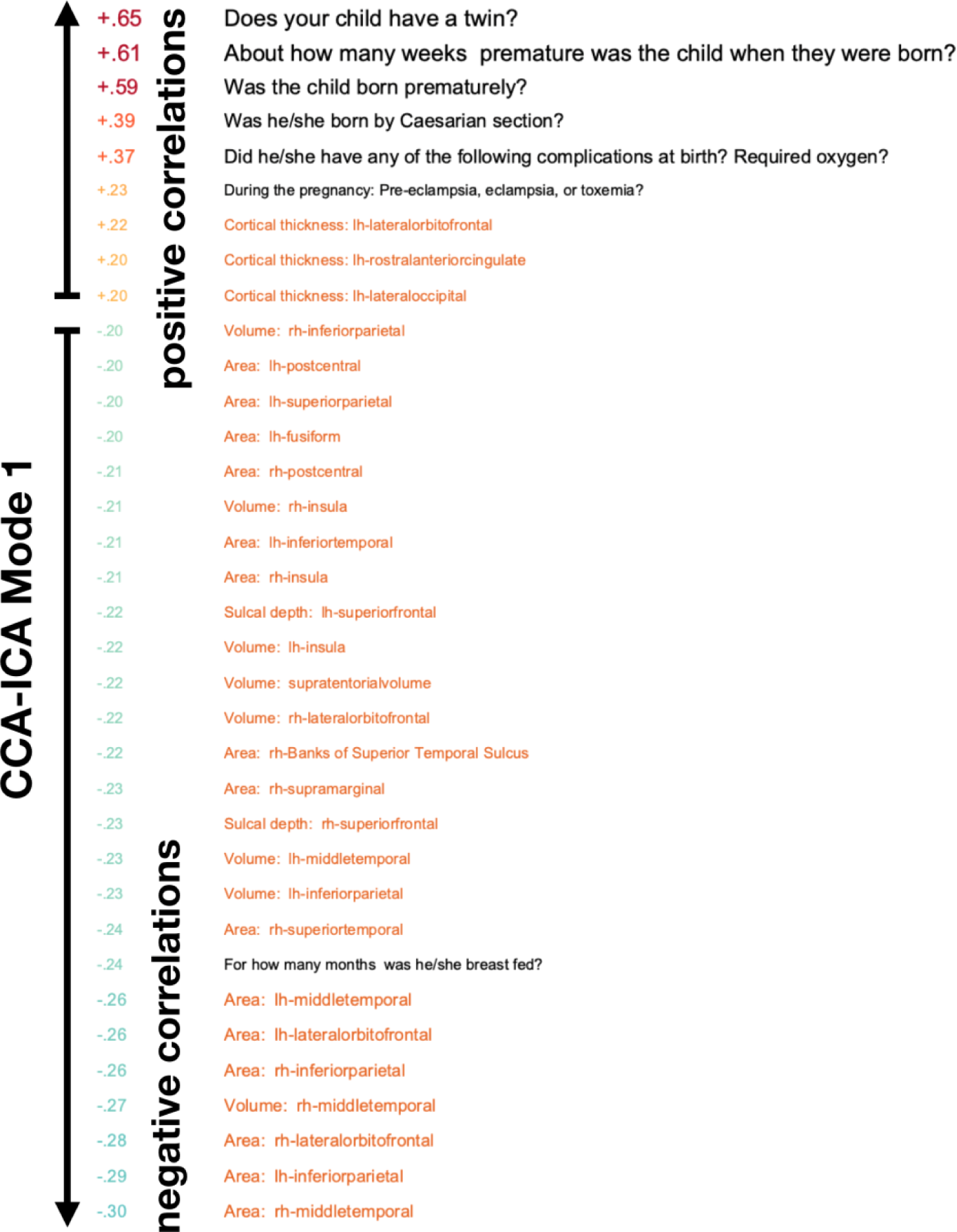
Mode 1 links obstetric and perinatal complications to cortical area, volume and thickness. Numbers on the left are correlations between each participant measure and MRI feature with Mode 1 CCA-ICA subject weights. Hot/cold colors represent positive/negative correlations, respectively. Text on the right represent the participant measures (black) and imaging features (orange). The top 35 items are shown, for a full list of all measures and their respective correlations see Supplementary Table 3

Mode 2 captures a pattern of economic deprivation and poverty, with the highest loading measures being related to the area deprivation index (ADI), such as parent unemployment, neighborhood median household income, income disparity and violence (**Fig. 3**). The mode links these measures to lower maternal age at child birth, lower parent education level, unplanned pregnancy, shorter duration of breast feeding, higher number of half-siblings, higher levels of religiosity, and the child having less nightly sleep hours on average, lower grades in school and worse performance on cognitive tests, jointly forming a dimension of socio-cognitive stratification. This dimension is associated with lower cortical thickness, area and volume, with total volume, lateral occipital cortical volumes and thickness, and bilateral lingual thickness among the highest-loading imaging features.

**Fig. 3:**
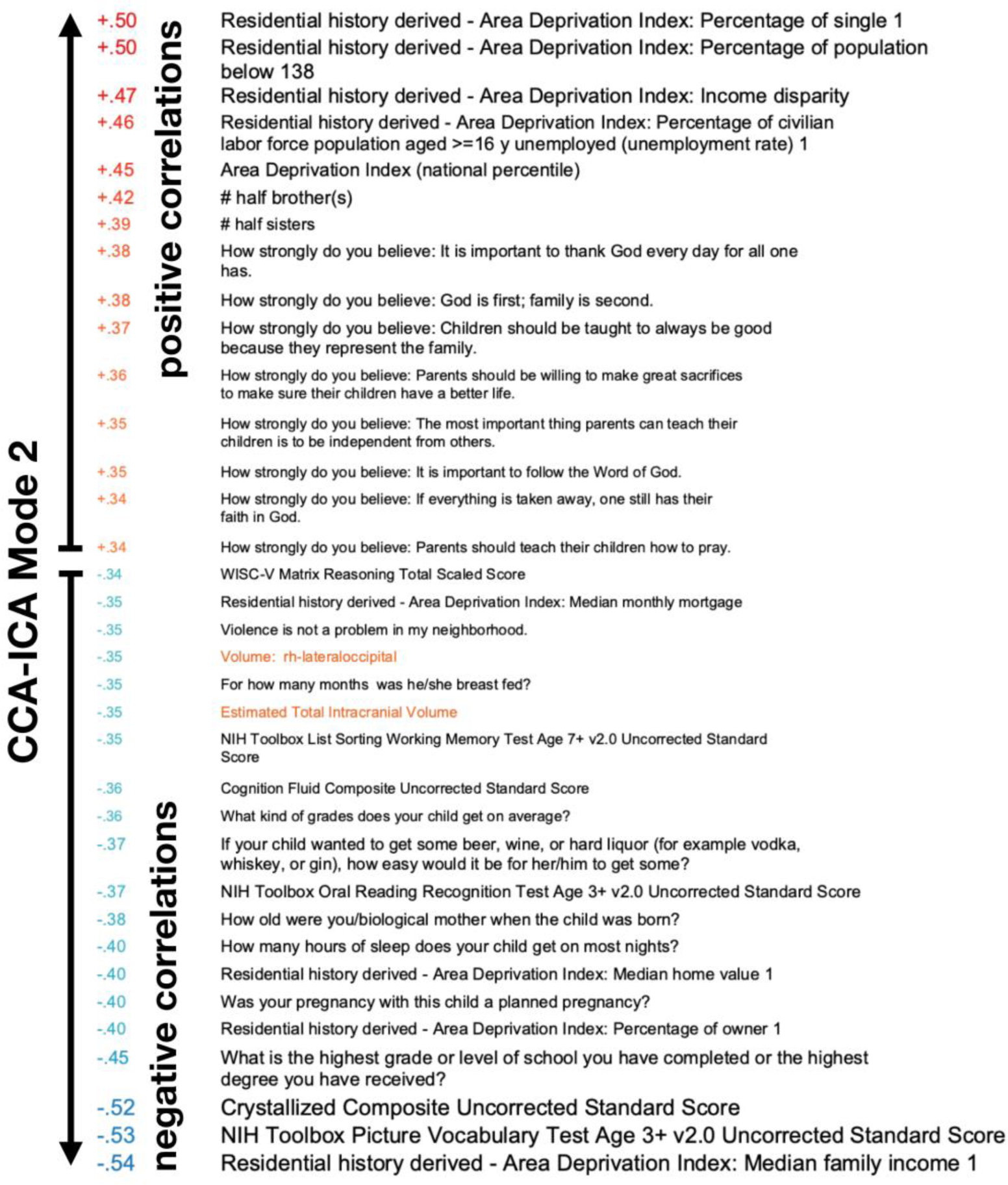
Variables with the highest correlations with CCA-ICA Mode 2 subject weights are shown, forming a dimension of socio-cognitive stratification, which is associated with total intracranial volume and regional cortical volumes and thickness. Numbers on the left are correlations, hot/cold colors represent positive/ negative correlations, respectively, arrows indicate increasing positive/negative correlations. Text on the right represents the behavioral measures (black) and imaging features (orange). The top 35 items are shown, for a full list of all measures and associated correlations see Supplementary Table 4.

Mode 3 captures a links higher air particle matter (PM^2.5^) and area deprivation to lower population density, lower levels of NO2, lower neighborhood walkability, lower home value and rent, but higher home ownership percentage, higher number of half-siblings, and living in a state which has not legalized marijuana for medical use (as of 2016). The mode (**Fig. 4, Supplementary Table 5**) is further associated with reporting emerging signs of puberty such as body hair and to lower area and volumes across the cortex, as well as with white matter indices such as fractional anisotropy (FA) radial diffusivity (RD) neurite density (ND) and tract volumes, with the highest loading measures being related to the parahippocampal cingulum, the uncinate fasciculus, and corpus callosum.

**Fig. 4:**
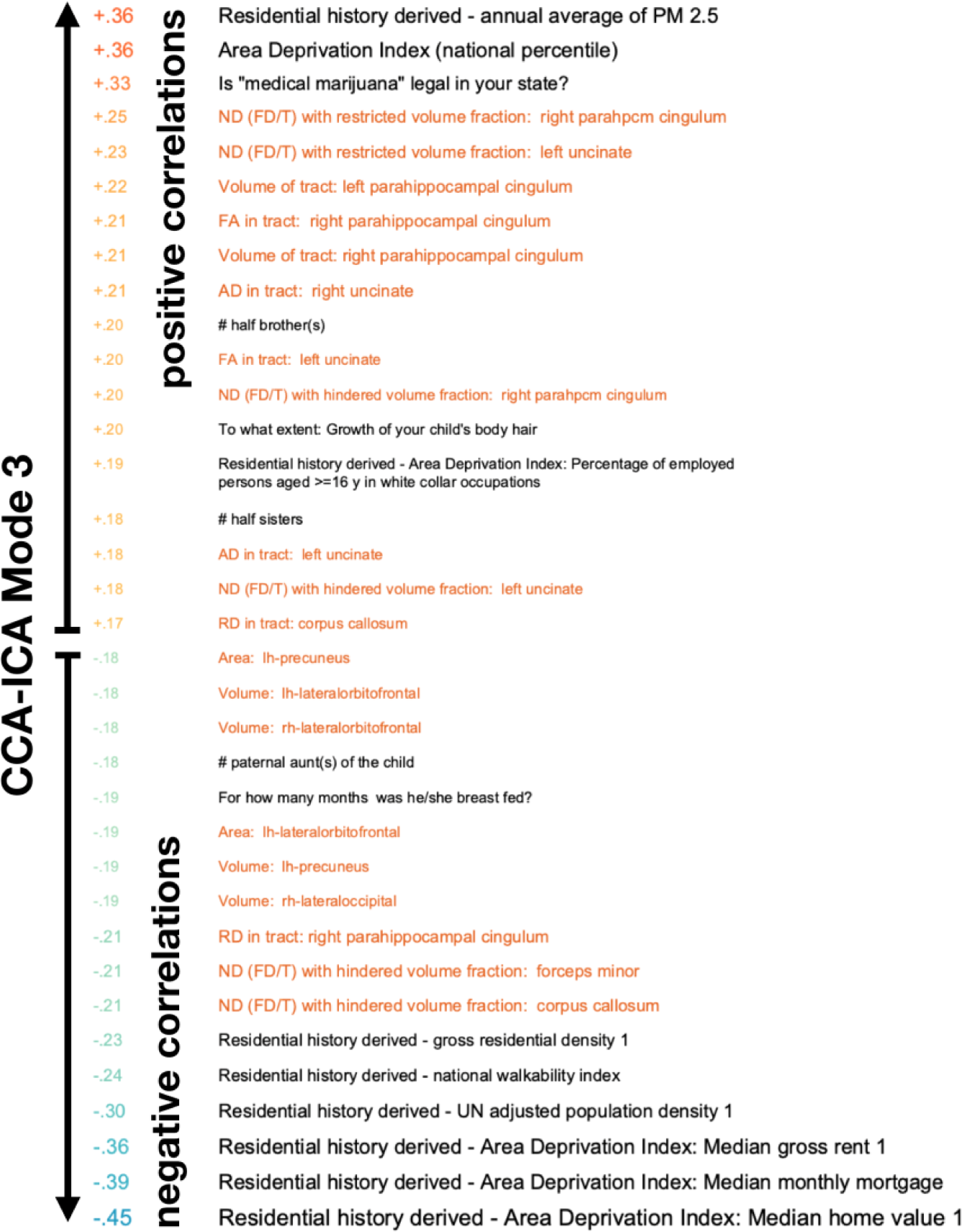
The variables with the highest correlations with CCA-ICA Mode 3 subject weights, linking air pollution, area deprivation, walkability and population density to brain white matter indices. Numbers on the left are correlations, hot/cold colors represent positive/negative correlations, respectively, arrows indicate increasing positive/negative correlations. Text on the right represents the behavioral measures (black) and imaging features (orange). The top 35 items are shown, for a full list of all measures and associated correlations see Supplementary Table 5.

## Discussion

Adolescence is a transition period between childhood to adulthood, associated with heightened sensitivity to the social and cultural environment^12^. While for most individuals the transition results in successful acquirement of skills and coping strategies required for adulthood and subsequent independence from caregivers, it also coincides with increased risk for mental health issues and psychological madadjustment^13^. Research addressing the social, economic and environmental conditions affecting adolescent development, facilitating health and leading to fulfilling adult lives is therefore critical. Here we discuss three modes of population co-variation, each linking behavioral, clinical, psychosocial, socioeconomic and demographical measures to neuroimaging in 7,577 children aged 9-11 years.

The first mode links obstetric complications and early life factors such as duration of breast feeding and motor development, with cognitive ability, cortical surface area, thickness and volume in late childhood. Obstetric complications increase the risk of later cognitive deficits and mental disorders^25^. The present results support that children with a history of obstetric and perinatal complications show delayed brain development and are consistent with reports associating birth weight with cortical area and brain volume in childhood and adolescence^26^, underscoring the importance of taking perinatal factors into account when studying child and adolescent brain development.

The second mode captures a socio-cognitive stratification pattern associated with brain volume and regional measures of cortical thickness, area and volume. Conceptually, the mode shares similarities with a positive-negative population mode linked to brain functional connectivity^1^ and structure^15^ in adults. The mode links several positive and negative life-events and environmental circumstances, with the highest loading factors being related to socioeconomic status, such as poverty, parent unemployment and education level. It further captures several factors known to be related to social deprivation, such as degree of family-planning and early pregnancies, neighborhood level of violence and level of religious beliefs. These constitute important environmental conditions for neurodevelopment that these children receive from their parents, their community and society at large. Consistent with the literature on the effect of social deprivation on child development, this mode is also associated with less sleep, worse school performance and lower cognitive ability. Cognitive ability is moderately heritable in childhood and adolesence^9, 27, 28^, and this is likely partly explaining the association between child cognition, academic performance and SES^29^. However, the effects of poverty, low socioeconomic status and early life adversity on brain and cognitive development^18, 30^ also underscore the role and importance of social policies aimed at reducing disparities^31^ that put some children at a disadvantage, often with life-long consequences for opportunities, mental and physical health and quality of life. The neurotypical developmental trajectory at this age is characterized by apparent cortical thinning, likely partly reflecting synaptic pruning^32^ and myelination^33^. Thicker cortex with higher SES is consistent with reports of accelerated brain maturation in children from low-SES families^34, 35^. Indeed, across species and in humans, early life adversity is associated with accelerated maturation of neural systems, possibly at a cost of increased risk for later mental health problems^36^.

The third mode reflects an inverse association between particulate matter air pollution (PM^2.5^) and area deprivation on one side, and home value, walkability, and population density on the other. Specific geographical information for this mode cannot be discerned, since the ABCD does not provide geographical data about its participants, however the this mode fits a known socio-economic settlement pattern: low-pollution, high-walkability “sweet-spot” neighborhoods in urban areas are typically skewed toward higher-SES households, contrasted with high-pollution, low-walkability “sour-spot” neighborhoods associated with lower income^37^. Exposure to PM^2.5,^ is associated with adverse health outcomes and disproportionally affects lower-income households^38^. Interestingly, this pattern was associated with the legal status of medical marijuana (as of 2016), possibly indicative of geographical differences for this pattern across the US states. Here we document an association with cortical area and volumes, as well as associations with diffusion properties of brain white matter pathways, in particular the parahippocampal cingulum, uncinate fasciculus, corpus callosum and forceps minor.

While these population level patterns are highly interesting, the cross-sectional and the non-experimental design warrant caution. People and their brains, genes and environments are not varying randomly, but are highly correlated^39^, likely along multiple dimensions, which complicates causal and mechanistic inference. This is especially relevant for population-based neuroimaging, in which subtle confounds can induce spurious associations^4^. These general caveats notwithstanding, these valuable resources represent an unprecedented opportunity to reveal co-varying patterns of socio-demographics, cognitive abilities, mental health, and brain imaging data, beyond simple bivariate associations, which are potentially highly informative of the biology, psychology and sociology of childhood and adolescent brain development and psychological adaptation.

In contrast to the standard regression approach which models one outcome-variable at a time and typically includes only a few covariates the combined multivariate approach employed here considers the full pattern of co-variability between variables. Our approach is therefore well suited for capturing population patterns by maximizing statistical power. However, it does not allow for interpretation of specific associations between pairs of variables. Overfitting can be a challenge with multivariate approaches, in particular in small samples and for complex models^40^. Currently there are no comparable samples to ABCD in which to independently assess the generalizability of the results. However, the current results were obtained in a large sample, using data reduction as well as 10-fold cross-validation with all relevant analysis steps performed within the cross-validation and permutation loop, to avoid over-fitting and assess generalizability. All the patterns are purely correlational and also treated analytically and reported as such. It is also entirely possible, and highly probable, that these patterns are further correlated with other important phenomena not measured or included in the current analysis. The current approach also effectively captures differential patterns involving the same measures. For example, higher cortical thickness, indicative of delayed maturation, is independently associated both with socio-cognitive stratification, higher cognitive ability and SES, as well as with obstetric and perinatal complications, lower cognitive ability, and delayed speech and motor development. Another example is duration of breast feeding, which where independently associated both with obstetric complications as well as with socio-cognitive stratification, associated with differential patterns of brain differences. These independent and co-existing associations with brain structure emphasize the importance of multidimensional considerations for understanding child and adolescent neurodevelopment and support that political priorities and decisions aiming to improve health outcomes and adaptation during transformative life phases should be based on interdisciplinary perspectives integrating social, psychological and biological sciences^41^.

## Acknowledgements

D.A. is funded by the South- Eastern Norway Regional Health Authority (2019107). T.K. is funded by the Research Council of Norway (276082). A.F.M. gratefully acknowledges support from the Dutch Organisation for Scientific Research via a Vernieuwingsimpuls VIDI fellowship (016.156.415) and a Wellcome Trust Innovator award (215698/Z/19/Z). S.M.S is funded by a Wellcome Trust grant (203139/Z/16/Z). L.T.W. is funded by the European Research Council under the European Union’s Horizon 2020 research and innovation program (ERC Starting Grant 802998), the Research Council of Norway (249795), the South-East Norway Regional Health Authority (2019101), and the Department of Psychology, University of Oslo.

## Supplementary information

### 1. Supplementary Figures

**Supplementary Fig. 1:**
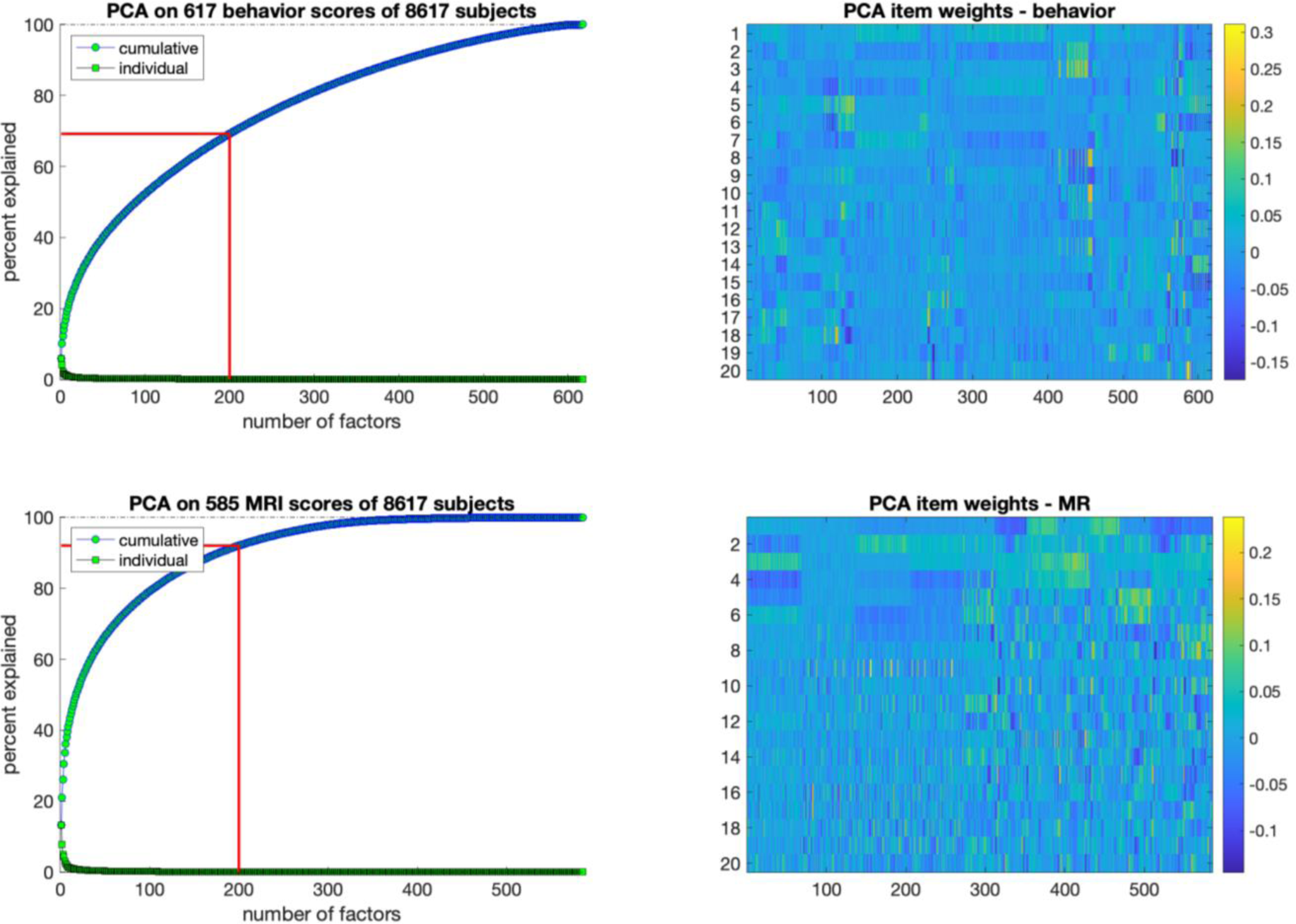
PCA-decompositions. Top and bottom panels show behavior/clinical/demographics, and imaging phenotypes, respectively. Left panels: Cumulative explained variance as a function of included eigenvalues/components. Red line demarks the 200 included PCA-components. Right panels: PCA-item weights for the first 20 components.

**Supplementary Fig. 2:**
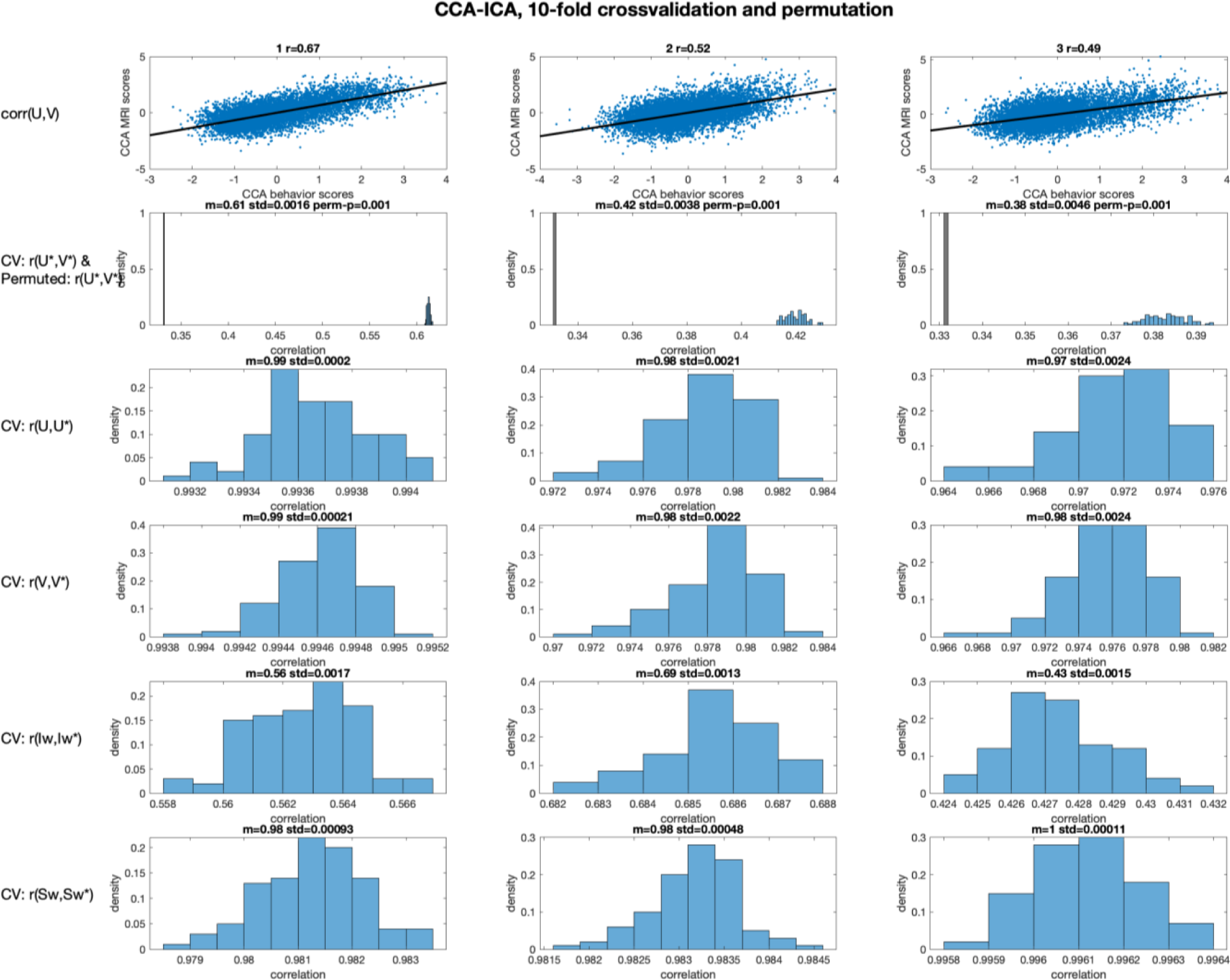
The four columns show results for CCA and CCA-ICA modes 1-4, respectively. **Row 1** shows the scatter-plots of behavioral and imaging scores from the in-sample CCA, and the title shows the canonical correlation for the full in-sample analysis (100% of data). **Row 2** Shown in blue is the distribution of out-of-sample canonical correlation values across 100 iterations of a 10-fold cross-validation procedure. Each fold (10% of participants) was kept out once, while estimating PCA and CCA using the remaining folds (90% of participants). Each of the 100 correlations plotted here is the mean canonical correlation between CCA weights derived for kept-out participants across the 10 folds in each of the 100 iterations. Title displays the mean out-of-sample canonical correlations, along with the standard deviation. The null-distribution generated by the permutation procedure is shown in black in the same plot: we ran 1000 iterations of the 10-fold cross-validation procedure, randomizing the rows (participants) of the imaging feature-matrix for each run. To correct for familywise error (FWE) we collected the largest canonical correlation (i.e. the value for mode 1) to form the null-distribution, and used this to calculate p-values using the mean canonical correlation for kept-out participants from the cross-validation procedure. **Row 3** shows the correlations between CCA behavioral measure-weights derived during cross-validation for folds kept out of the estimation, with those from the full analysis. **Row 4** shows the correlations between CCA imaging feature-weights derived during cross-validation for folds kept out of the estimation, with those from the full analysis. **Row 5** shows the correlations between ICA behavioral-measure and imaging-feature weights derived during cross-validation for folds kept out of the estimation, with those from the full analysis. **Row 6** shows the correlations between ICA subject weights derived during cross-validation for folds kept out of the estimation, with those from the full analysis. Y-axis in all histograms represent densities.

**Supplementary Fig. 3:**
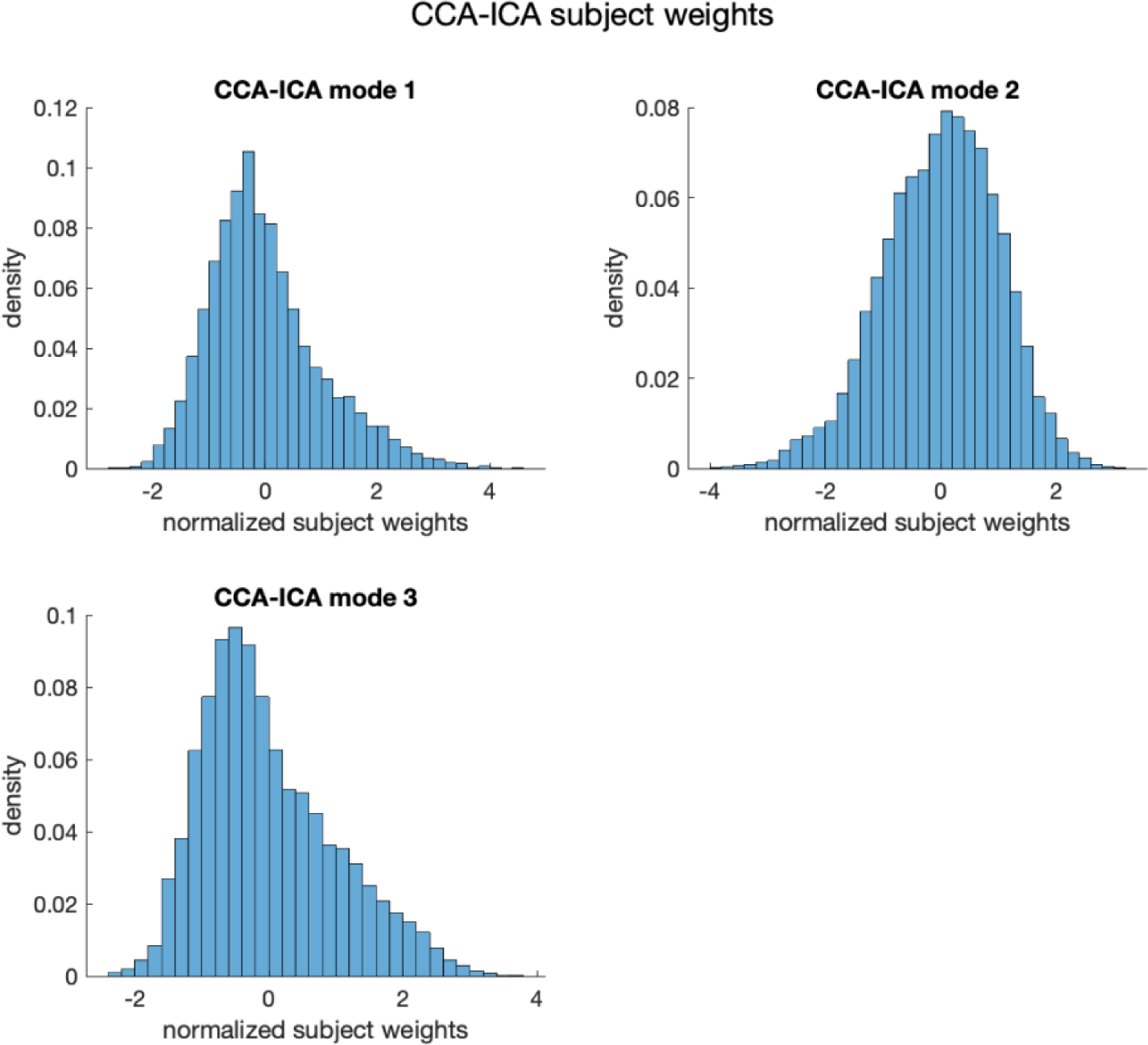
Histograms of CCA-ICA subject weights for the three significant modes.

**Supplementary Fig. 4:**
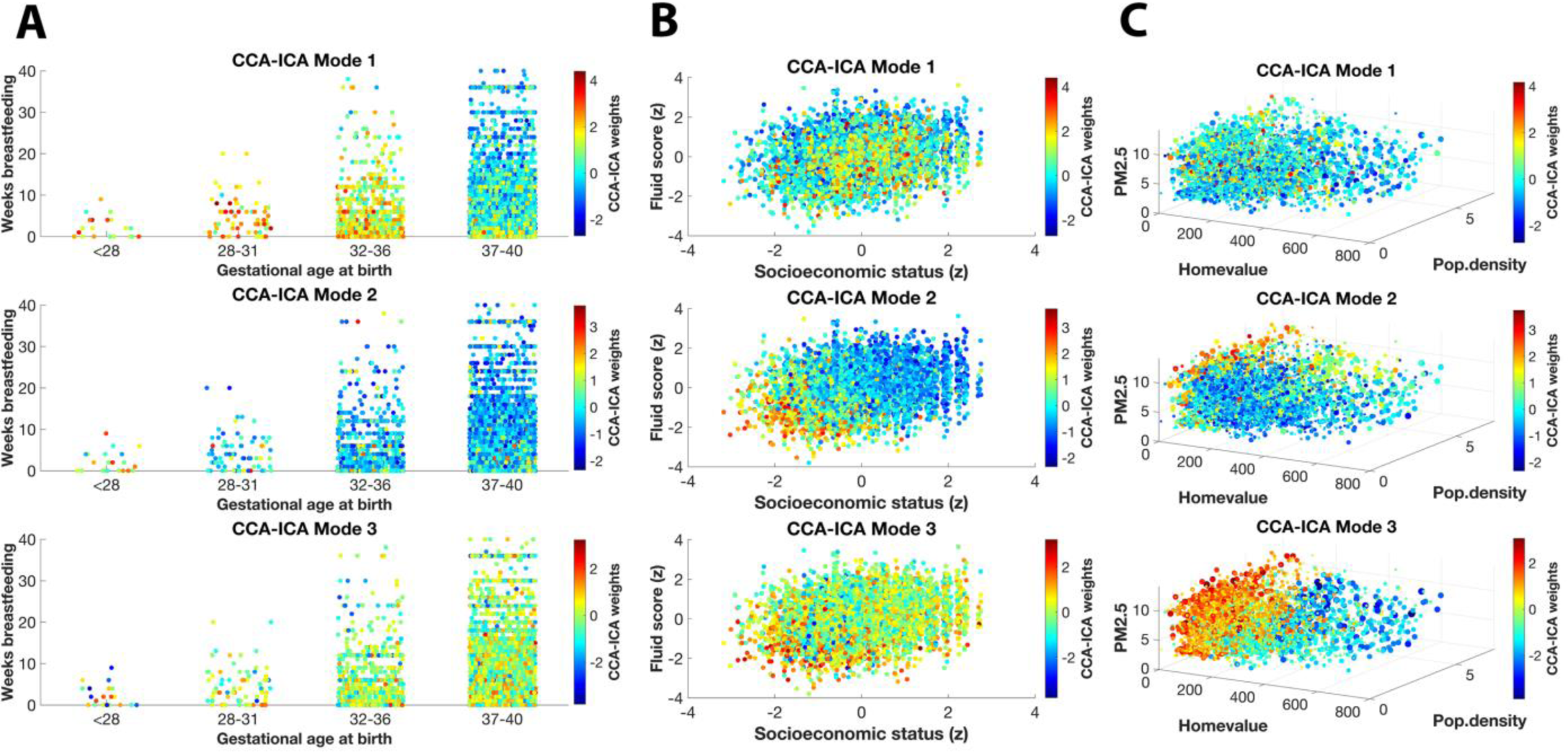
Plots visualizing associations among highest loading variables on the three CCA-ICA modes color-coded by the subject loadings for each mode. **A**: The groups on the X-axis represent gestational age at birth, the Y-axis shows the duration of breastfeeding, which are the highest loading items on Mode 1 (top row). **B:** The X-axis represents normalized SES-scores (based on parent education, parent income, and items related to whether the family can afford food, mortgage payments, phone bill, electricity and medical/dental services), and the Y-axis the child fluid composite score from the ABCD cognitive test battery, which are the highest loading items on Mode 2 (middle row). **C:** The X and Y and Z-axis show home-value, population density and particle matter pollution (PM^25^), respectively, the highest loading items on Mode 3 (bottom row).

#### Supplementary Fig. 5 - 7 – Modes 1 - 3 by sex

**Supplementary Fig. 5:**
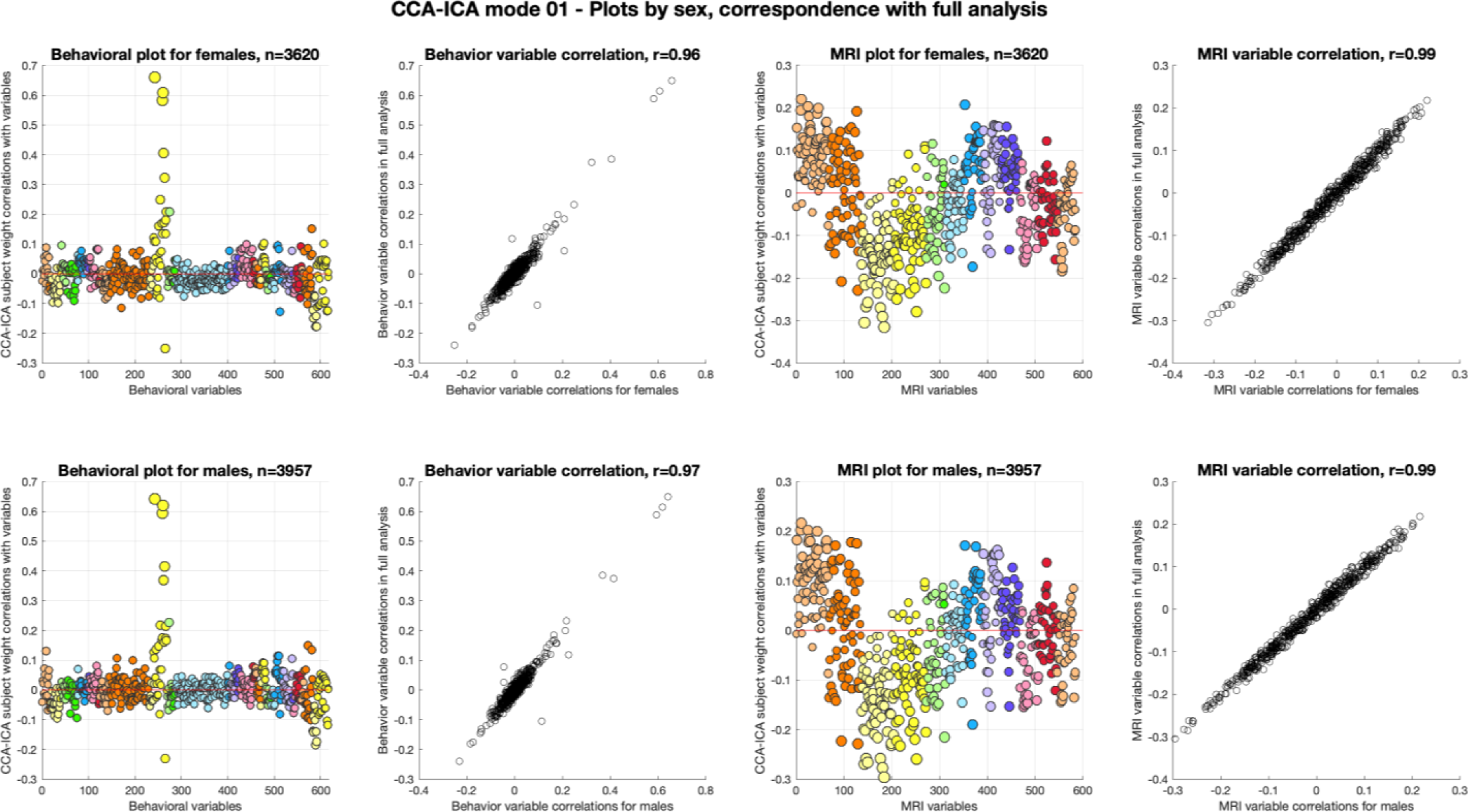
Colored plots show the correlation between the CCA-ICA mode 1 subject weights and the adjusted original behavioral and MRI data, plotted separately for females (F, top panels) and males (M, bottom panels), while scatter-plots shows the r-values for correlations between all CCA-ICA subject weights and the adjusted original behavioral and MRI data, plotted against the r-values for sex-specific CCA-ICA subject weight correlations.

**Supplementary Fig. 6:**
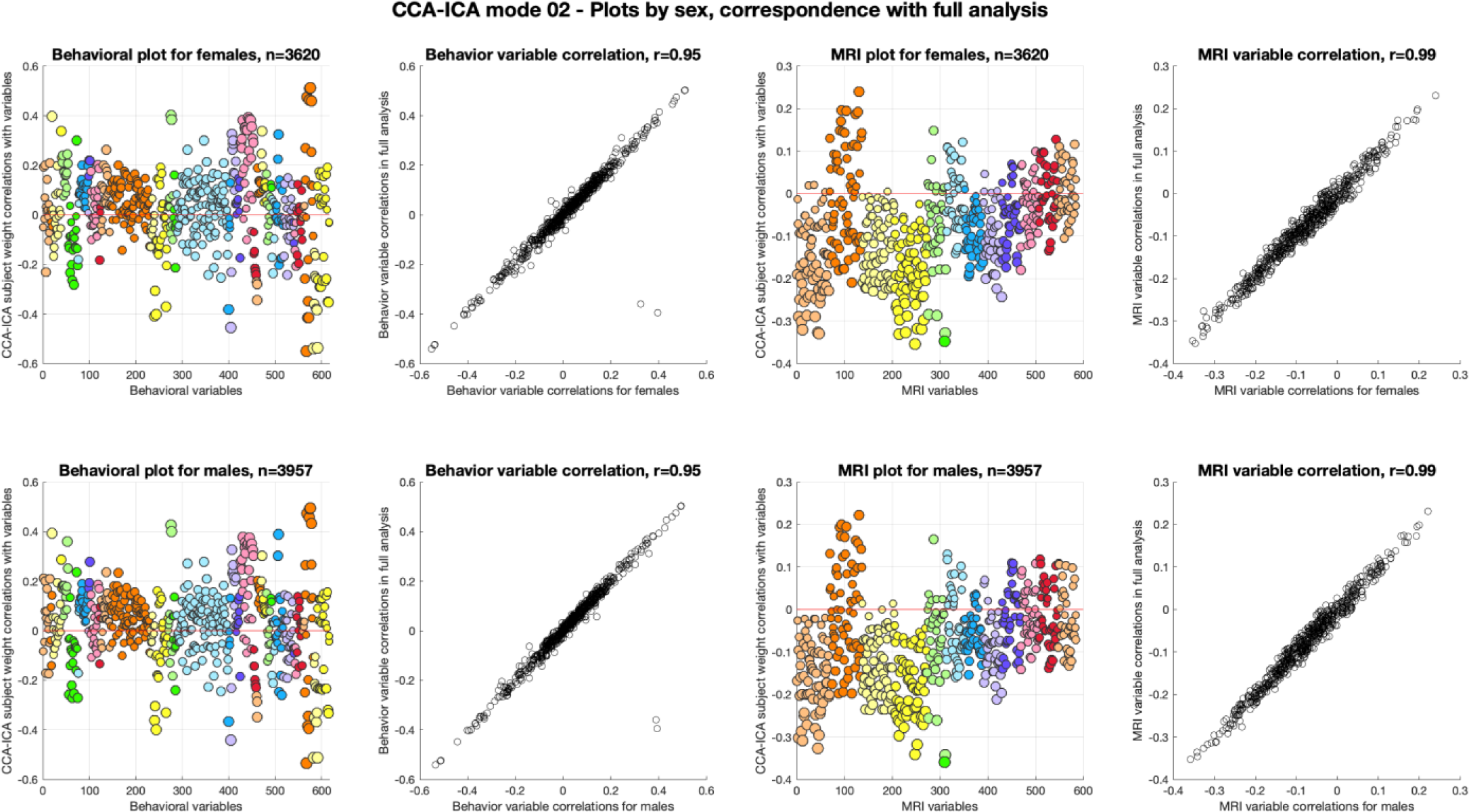
Colored plots show the correlation between the CCA-ICA mode 2 subject weights and the adjusted original behavioral and MRI data, plotted separately for females (F, top panels) and males (M, bottom panels), while scatter-plots shows the r-values for correlations between all CCA-ICA subject weights and the adjusted original behavioral and MRI data, plotted against the r-values for sex-specific CCA-ICA subject weight correlations.

**Supplementary Fig. 7:**
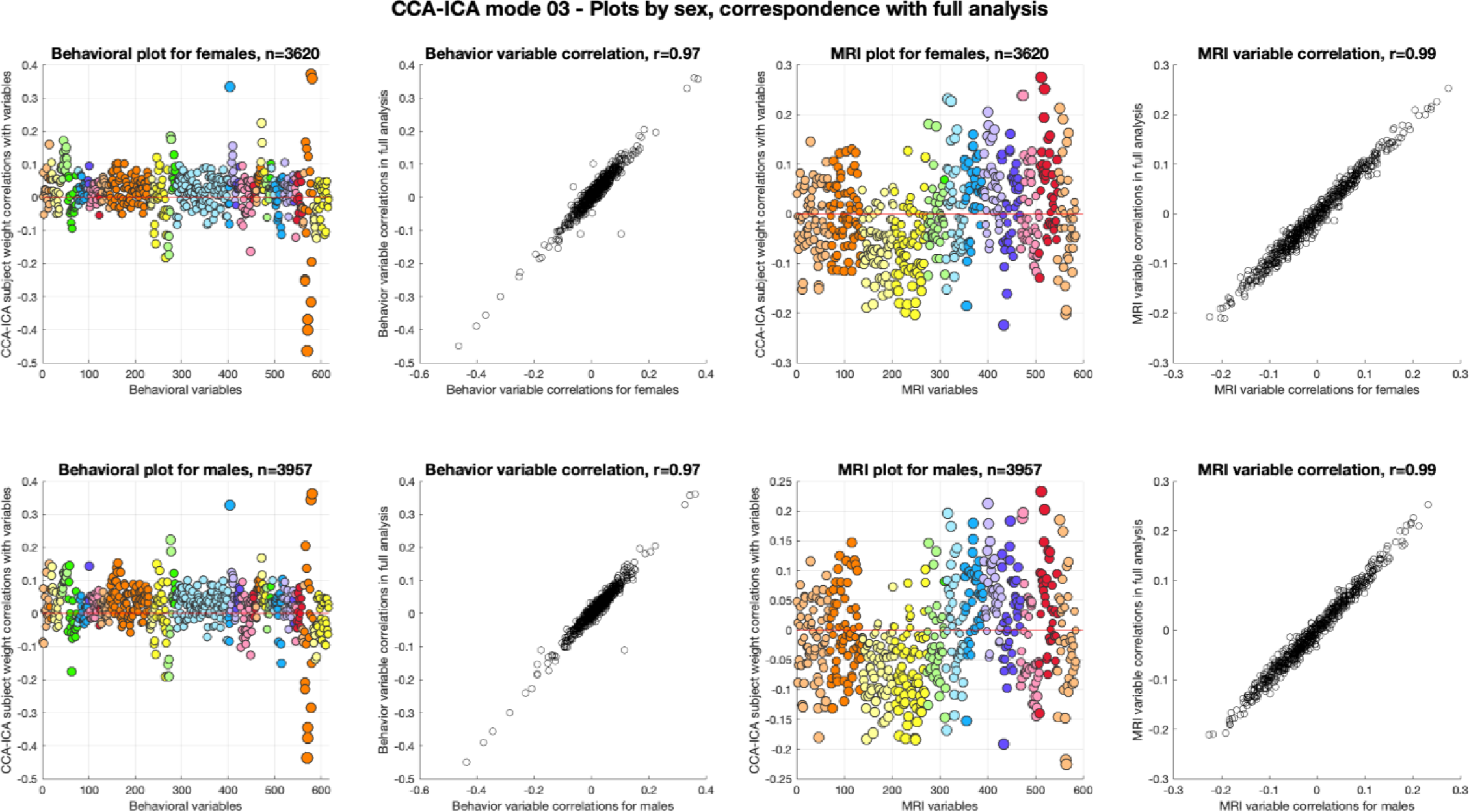
Colored plots show the correlation between the CCA-ICA mode 3 subject weights and the adjusted original behavioral and MRI data, plotted separately for females (F, top panels) and males (M, bottom panels), while scatter-plots shows the r-values for correlations between all CCA-ICA subject weights and the adjusted original behavioral and MRI data, plotted against the r-values for sex-specific CCA-ICA subject weight correlations.

#### Supplementary Fig. 8-10: Mode 1 – 3 by ethnicity/race

**Supplementary Fig. 8:**
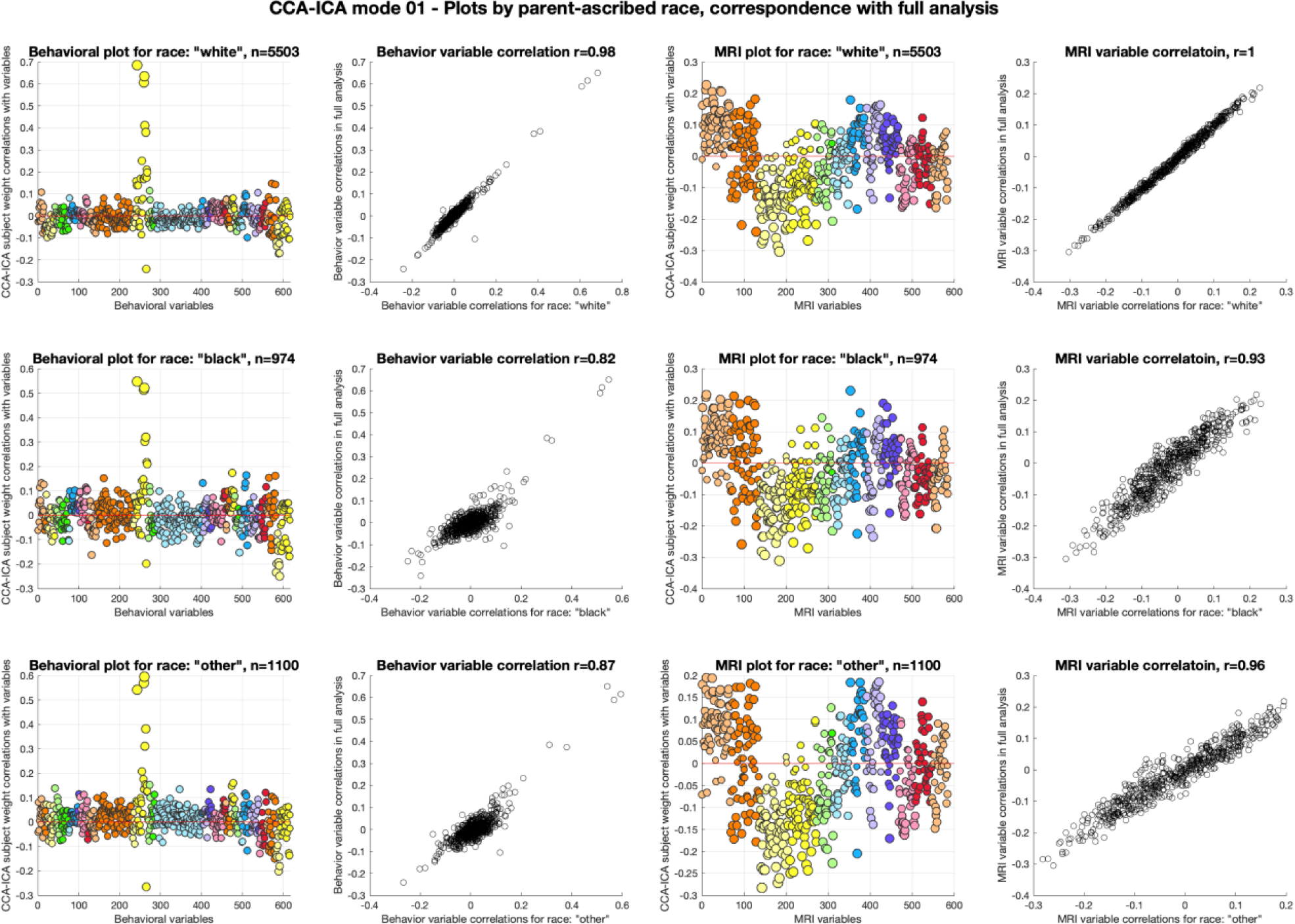
Colored plots show the correlation between the CCA-ICA mode 1 subject weights and the adjusted original behavioral and MRI data, plotted separately for parent designated race (“white” in top panels, “black” in middle panels, “other” in bottom panels, other possible categories constituted less than 5% of the sample), while scatter-plots shows the r-values for correlations between all CCA-ICA subject weights and the adjusted original behavioral and MRI data plotted against the r-values for parent-designated race-specific CCA-ICA subject weight correlations.

**Supplementary Fig. 9:**
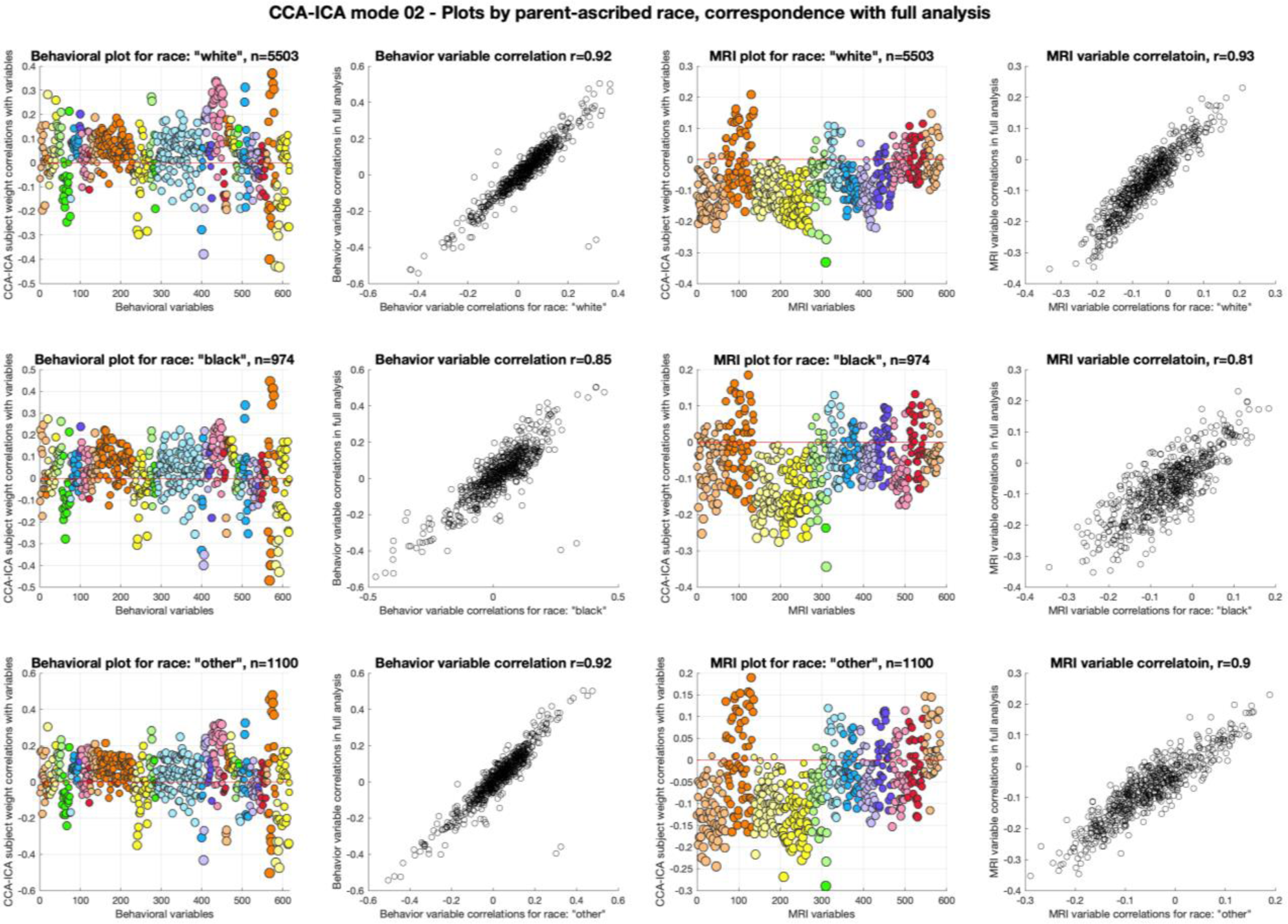
Colored plots show the correlation between the CCA-ICA mode 2 subject weights and the adjusted original behavioral and MRI data, plotted separately for parent designated race (“white” in top panels, “black” in middle panels, “other” in bottom panels, other possible categories constituted less than 5% of the sample), while scatter-plots shows the r-values for correlations between all CCA-ICA subject weights and the adjusted original behavioral and MRI data plotted against the r-values for parent-designated race-specific CCA-ICA subject weight correlations.

**Supplementary Fig. 10:**
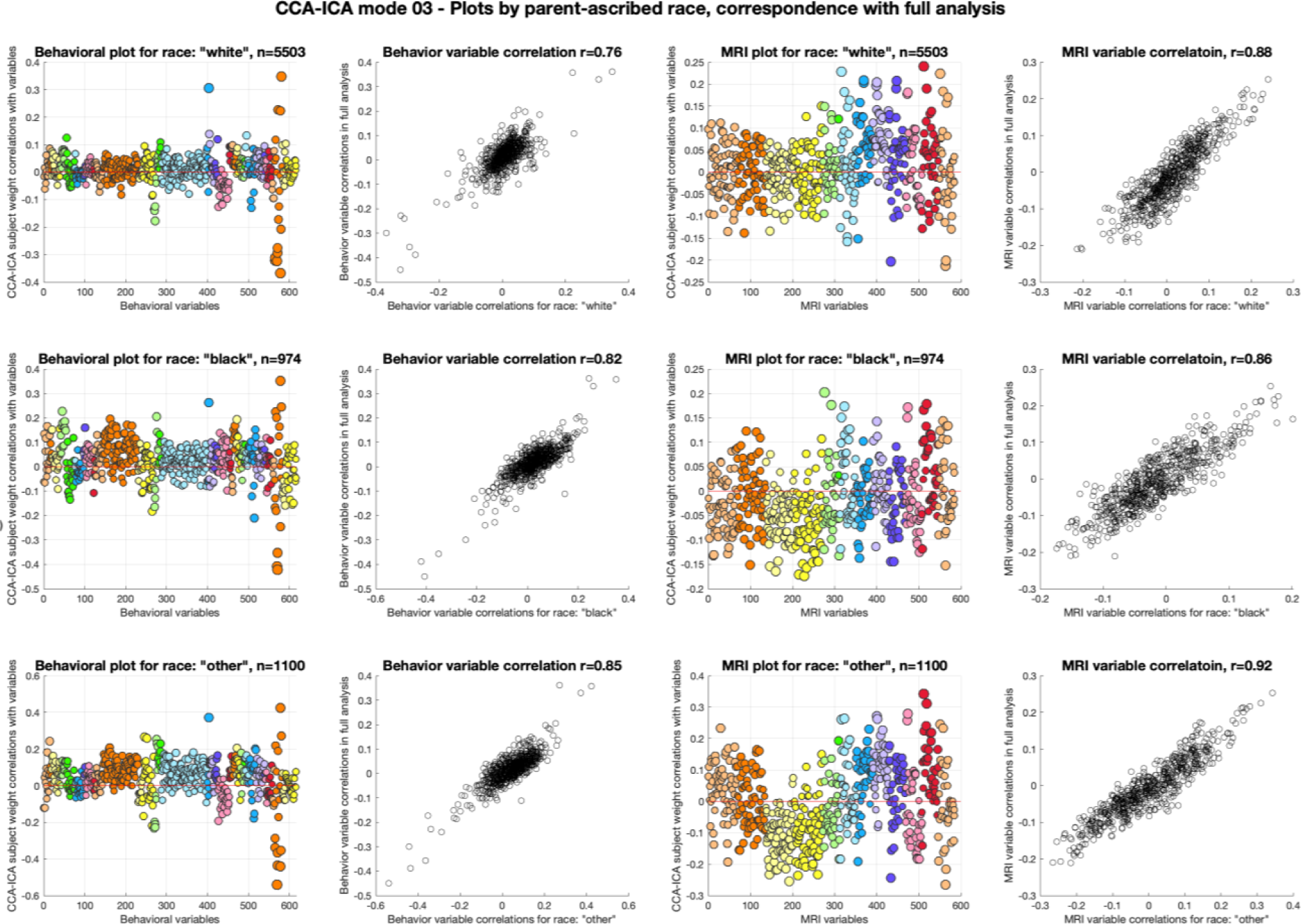
Colored plots show the correlation between the CCA-ICA mode 3 subject weights and the adjusted original behavioral and MRI data, plotted separately for parent designated race (“white” in top panels, “black” in middle panels, “other” in bottom panels, other possible categories constituted less than 5% of the sample), while scatter-plots shows the r-values for correlations between all CCA-ICA subject weights and the adjusted original behavioral and MRI data plotted against the r-values for parent-designated race-specific CCA-ICA subject weight correlations.

#### Supplementary Fig. 11-13: Mode 1 – 3 by site/scanner

**Supplementary Fig. 11:**
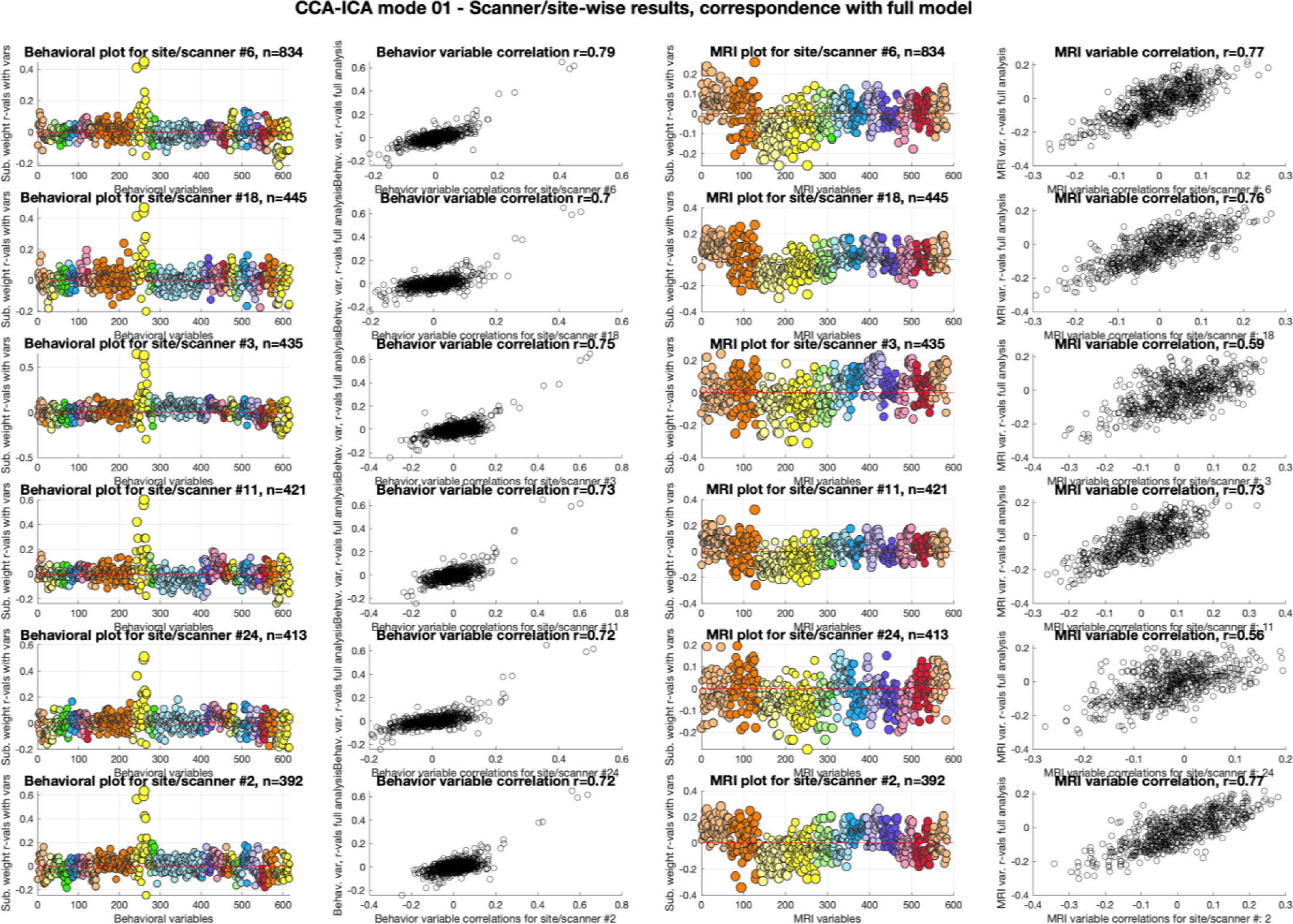
Colored plots show the correlation between the CCA-ICA mode 1 subject weights and the adjusted original behavioral and MRI data, plotted separately for the sites/scanner constituting >= 5% of the total sample, in descending order, while scatter-plots shows the r-values for correlations between all CCA-ICA subject weights and the adjusted original behavioral and MRI data plotted against the r-values for scanner/site-specific CCA-ICA subject weight correlations.

**Supplementary Fig. 12:**
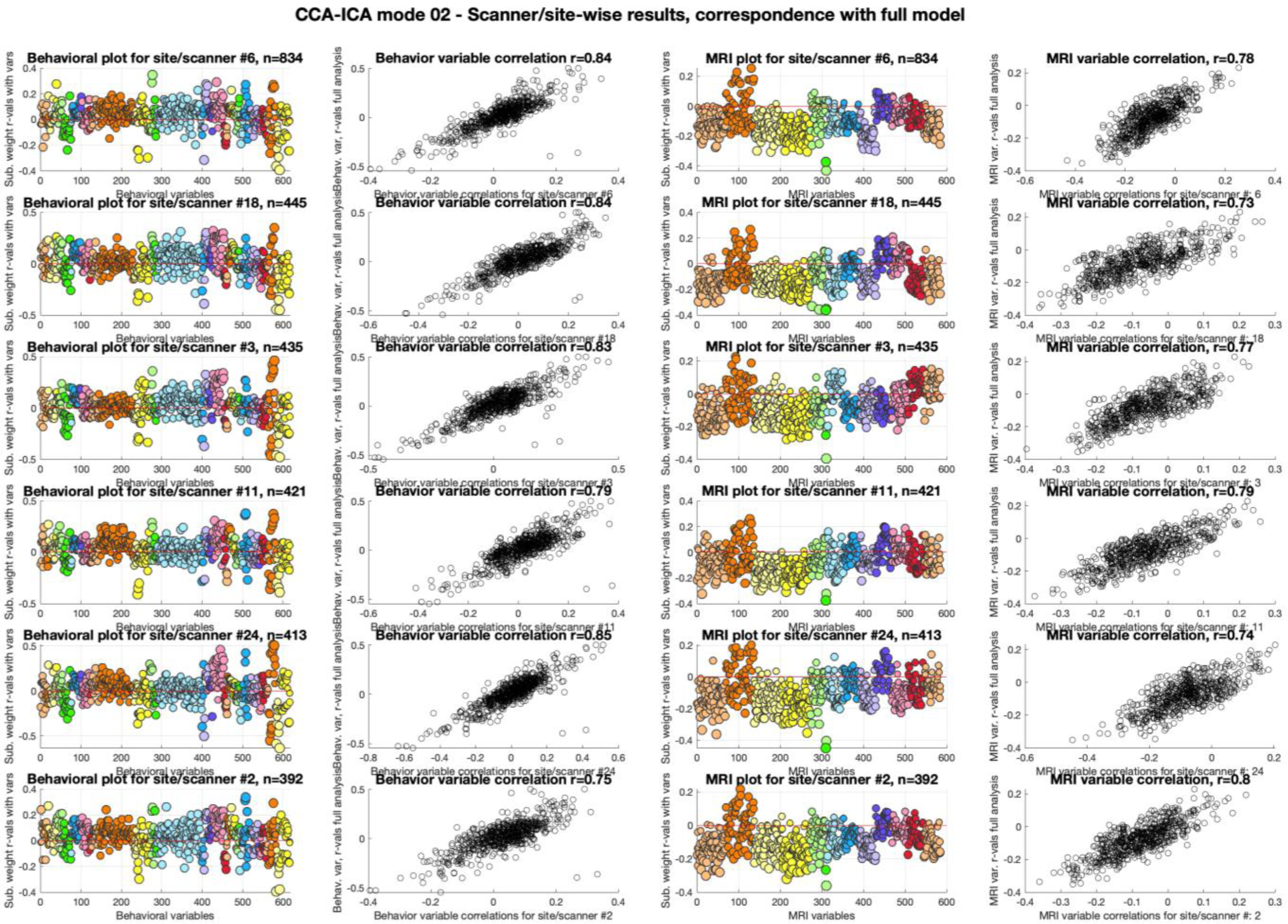
Colored plots show the correlation between the CCA-ICA mode 2 subject weights and the adjusted original behavioral and MRI data, plotted separately for the sites/scanner constituting >= 5% of the total sample, in descending order, while scatter-plots shows the r-values for correlations between all CCA-ICA subject weights and the adjusted original behavioral and MRI data plotted against the r-values for scanner/site-specific CCA-ICA subject weight correlations.

**Supplementary Fig. 13:**
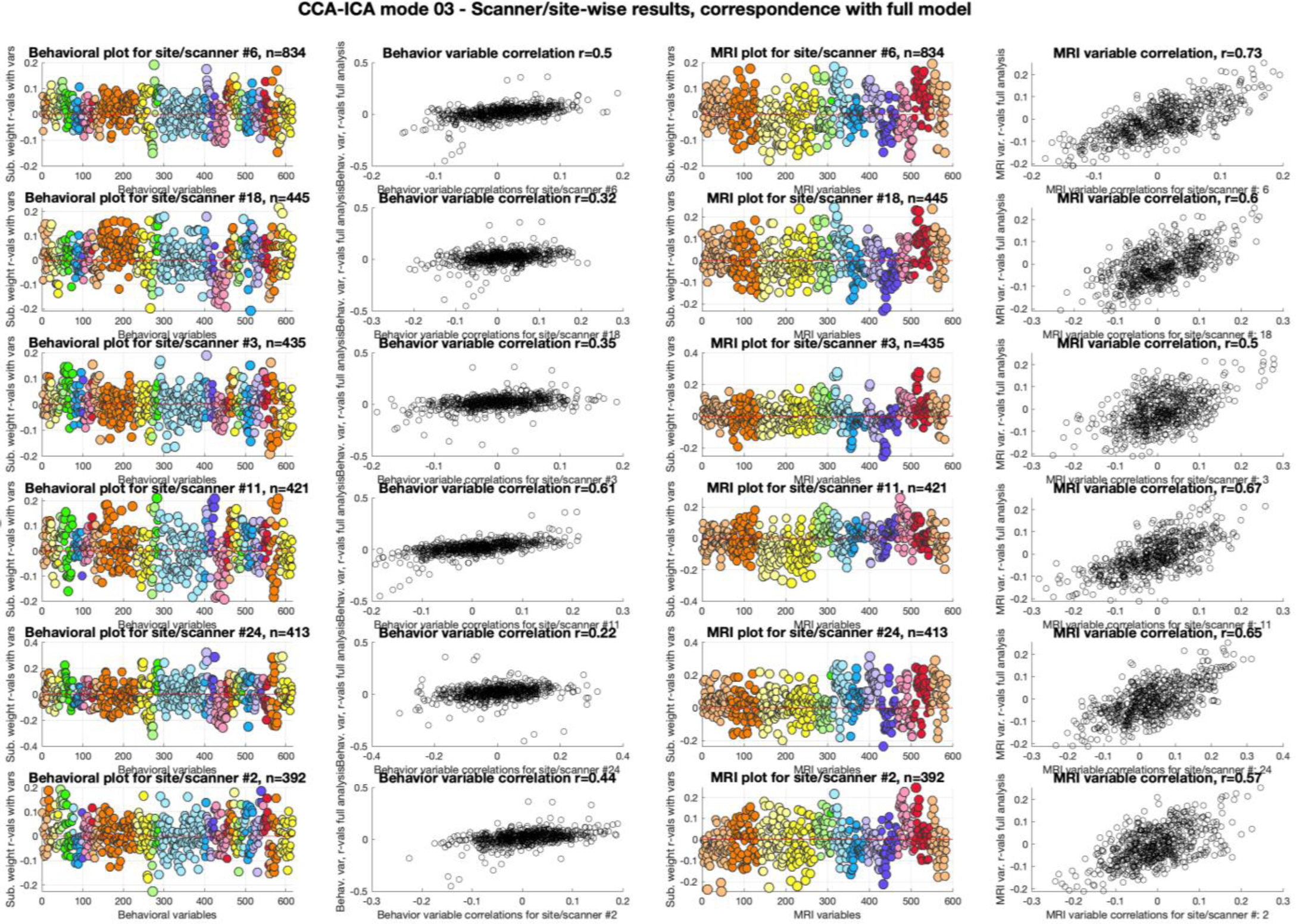
Colored plots show the correlation between the CCA-ICA mode 3 subject weights and the adjusted original behavioral and MRI data, plotted separately for the sites/scanner constituting >= 5% of the total sample, in descending order, while scatter-plots shows the r-values for correlations between all CCA-ICA subject weights and the adjusted original behavioral and MRI data plotted against the r-values for scanner/site-specific CCA-ICA subject weight correlations.

### 2. Supplementary Tables

Tables available using this link to a OSF.io data repository (Center for Open Science) for the current paper.

**Supplementary Table 1:** Lists the included behavioral variable names, and their associated descriptions and instrument.

**Supplementary Table 2:** Lists the included MRI variable names, and their associated descriptions.

**Supplementary Tables 3-5:** List of all included variables, their description, ICA-weights and correlations with CCA-ICAs Mode 1-3 subject weights.

## References

1. Smith, S.M., et al. Nature neuroscience 18, 1565 (2015).

2. Casey, B.J., et al. Developmental Cognitive Neuroscience 32, 43–54 (2018).

3. Paulus, M.P. & Thompson, W.K. JAMA Psychiatry 76, 353–354 (2019).

4. Smith, S.M. & Nichols, T.E. Neuron 97, 263–268 (2018).

5. Finn, E.S., et al. Nature neuroscience 18, 1664 (2015).

6. Kaufmann, T., et al. Nature neuroscience 20, 513 (2017).

7. Kaufmann, T., et al. Nature neuroscience (2019).

8. Franke, K. & Gaser, C. Frontiers in Neurology 10, 789 (2019).

9. Alnæs, D., et al. JAMA Psychiatry (2018).

10. Watanabe, K., et al. Nature Genetics 51, 1339–1348 (2019).

11. Visscher, P.M., et al. The American Journal of Human Genetics 101, 5–22 (2017).

12. Blakemore, S.-J. & Mills, K.L. Annual review of psychology 65, 187–207 (2014).

13. Paus, T., Keshavan, M. & Giedd, J.N. Nature reviews. Neuroscience 9, 947–957 (2008).

14. Miller, K.L., et al. Nature neuroscience 19, 1523 (2016).

15. Llera, A., Wolfers, T., Mulders, P. & Beckmann, C.F. Elife 8, e44443 (2019).

16. Bor, J., Cohen, G.H. & Galea, S. The Lancet 389, 1475–1490 (2017).

17. Hair, N.L., Hanson, J.L., Wolfe, B.L. & Pollak, S.D. JAMA Pediatrics 169, 822–829 (2015).

18. Noble, K.G., et al. Nature neuroscience 18, 773–778 (2015).

19. Leys, C., Ley, C., Klein, O., Bernard, P. & Licata, L. Journal of Experimental Social Psychology 49, 764–766 (2013).

20. Hagler, D.J., et al. NeuroImage, 116091 (2019).

21. Yang, J., Lee, S.H., Goddard, M.E. & Visscher, P.M. American Journal of Human Genetics 88, 76–82 (2011).

22. Krzanowski, W.J. Principles of multivariate analysis: a user’s perspective (Oxford University Press, Inc., 1988).

23. Winkler, A.M., Ridgway, G.R., Webster, M.A., Smith, S.M. & Nichols, T.E. NeuroImage 92, 381–397 (2014).

24. Hyvarinen, A. IEEE Transactions on Neural Networks 10, 626–634 (1999).

25. Mezquida, G., et al. The Journal of Nervous and Mental Disease 206 (2018).

26. Walhovd, K.B., et al. Proceedings of the National Academy of Sciences 109, 20089 (2012).

27. Allegrini, A.G., et al. Molecular psychiatry 24, 819–827 (2019).

28. Plomin, R. & Deary, I.J. Molecular psychiatry 20, 98–108 (2015).

29. Trzaskowski, M., et al. Intelligence 42, 83–88 (2014).

30. Johnson, S.B., Riis, J.L. & Noble, K.G. Pediatrics 137, e20153075 (2016).

31. Hermansen, A.S., Borgen, N.T. & Mastekaasa, A. European Sociological Review (2019).

32. Gilmore, J.H., Knickmeyer, R.C. & Gao, W. Nature Reviews Neuroscience 19, 123 (2018).

33. Natu, V.S., et al. Proceedings of the National Academy of Sciences 116, 20750 (2019).

34. LeWinn, K.Z., Sheridan, M.A., Keyes, K.M., Hamilton, A. & McLaughlin, K.A. Nature Communications 8, 874 (2017).

35. Piccolo, L.R., et al. PloS one 11, e0162511 (2016).

36. Callaghan, B.L. & Tottenham, N. Curr Opin Behav Sci 7, 76–81 (2016).

37. Marshall, J.D., Brauer, M. & Frank, L.D. Environ Health Perspect 117, 1752–1759 (2009).

38. Bowe, B., Xie, Y., Yan, Y. & Al-Aly, Z. JAMA Network Open 2, e1915834–e1915834 (2019).

39. Turkheimer, E. in Philosophy of behavioral biology. 43–64 (Springer Science, New York, NY, US, 2012).

40. Poldrack, R.A., Huckins, G. & Varoquaux, G. JAMA Psychiatry (2019).

41. Farah, M.J. Nature Reviews Neuroscience 19, 428–438 (2018).

